# Comparative genomics provides insight into the function of broad-host range sponge symbionts

**DOI:** 10.1101/2020.12.09.417808

**Authors:** Samantha C. Waterworth, Shirley Parker-Nance, Jason C. Kwan, Rosemary A. Dorrington

## Abstract

As the oldest extant metazoans, sponges (Phylum *Porifera*) have been forming symbiotic relationships with microbes that may date back as far as 700 million years. Most symbionts are conserved within a narrow host range and perform specialized functions. However, there are widely distributed bacterial taxa such as *Poribacteria, SAUL* and *Tethybacterales* that are found in a broad range of invertebrate hosts. Here, we added eleven new genomes to the *Tethybacterales* order, identified a novel family, and show that functional potential differs between the three *Tethybacterales* families. We compare the *Tethybacterales* with the well-characterized *Entoporibacteria* and show that these broad-host range, sponge-associated bacteria likely perform distinct functions within their hosts and that their respective phylogenies are incongruent with their host phylogenies. These results suggests that ancestors of these bacteria may have undergone multiple association events, rather than a single association event followed by co-evolution.

**IMPORTANCE:** Marine sponges often form symbiotic relationships with bacteria that fulfil a specific need within the sponge holobiont, and these symbionts are often conserved within a narrow range of related taxa. To date, there exist only three know bacterial taxa (*Entoporibacteria, SAUL* and *Tethybacterales*) that are globally distributed and found in a broad range of sponge hosts, and little is known about the latter two. Understanding what distinguishes these broad-host range symbionts from specialized symbionts will provide insight into the mechanisms by which sponges form these symbioses. We show that the functional potential of broad-host range symbionts is conserved at a family level and that these symbionts have been acquired several times over evolutionary history. This contrasts with specialized symbionts, where function is often a strain-specific trait and have co-evolved with their host following a single association event.

## INTRODUCTION

Sponges (Phylum *Porifera*) are the oldest living metazoans and they play an important role in maintaining the health of marine ecosystems(1). Sponges are remarkably efficient filter feeders, acquiring nutrients via phagocytosis of particulate matter, compromising mainly microbes, from the surrounding water(2). Since their emergence almost 700 million years ago, sponges have evolved close associations with microbial symbionts that provide services essential for the fitness and survival of the host in diverse ecological niches(3). These symbionts are involved in a diverse array of beneficial processes, including the cycling of nutrients(4, 5) such as nitrogen(5–10), sulfur(11, 12), phosphate(13, 14), the acquisition of carbon(15, 16), and a supply of vitamins(17–20) and amino acids(17). They can play a role in the host sponge life cycle, such as promoting larval settlement(21). In addition, some symbionts provide chemical defenses against predators and biofouling through the production of bioactive compounds(22–25). In turn, the sponge host can provide symbionts with nutrients and minerals, such as creatinine and ammonia as observed in *Cymbastela concentrica* sponges(8). In most cases, these specialized functions are performed by bacterial populations that are conserved within a given host taxon.

As filter-feeders, sponges encounter large quantities of bacteria and other microbes. How sponges are able to distinguish between prey bacteria and those of potential benefit to the sponge, and the establishment of symbiotic relationships, is still not well understood, but the structure and composition of bacterial lipopolysaccharide, peptidoglycan, or flagellin may aid the host sponge in distinguishing symbionts from prey(26). Sponge hosts encode an abundance of Nucleotide-binding domain and Leucine-rich Repeat (NLR) receptors, that recognize different microbial ligands and potentially allow for distinction between symbionts, pathogens, and prey(27). Additionally, it has recently been shown that phages produce ankyrins which modulate the sponge immune response and allow for colonization by bacteria(28).

Symbionts are often specific to their sponge host with enriched populations relative to the surrounding seawater(29). However, there are a small number of “cosmopolitan” symbionts that are ubiquitously distributed across phylogenetically distant sponge hosts. The *Poribacteria* and “*sponge-associated unclassified lineage*” (*SAUL*)(30) are examples of such cosmopolitan bacteria species. The phylum *Poribacteria* were thought to be exclusively found in sponges(31). However, the identification of thirteen putative *Poribacteria*-related metagenome-assembled genomes (MAGs) from ocean water samples(32), led to the reclassification and distinction of sponge-associated *Entoporibacteria* and free-living *Pelagiporibacteria* within the phylum(33). *Entoporibacteria* are associated with phylogenetically divergent sponge hosts in distant geographic locations, with no apparent correlations between their phylogeny and that of their sponge host or location(33, 34). Different *Poribacteria* phylotypes have been detected within the same sponge species(35). The *Entoporibacteria* carry several genes(36, 37) that encode enzymes responsible for the degradation of carbohydrates, and metabolism of sulfates and uronic acid(38–40), prompting the hypothesis that these bacteria may be involved in the breakdown of the proteoglycan host matrix(40). However, subsequent analyses of *Poribacteria* transcriptomes from the mesohyl of *Aplysina aerophoba* sponges showed that genes involved in carbohydrate metabolism were not highly expressed(39). Instead, there was a higher expression of genes involved in 1,2-propanediol degradation and import of vitamin B12, which together suggest that the bacterium may import vitamin B12 as a necessary cofactor for anaerobic 1,2-propanediol degradation and energy generation(39).

The *SAUL* bacteria belong to the larger taxon of candidate phylum *PAUC34f* and have been detected, although at low abundance, in several sponge species(41). Host-associated *SAUL* bacteria were likely acquired by eukaryotic hosts (sponges, corals, tunicates) at different evolutionary time points and are phylogenetically distinct from their planktonic relatives(41). Previous investigations into the only two SAUL bacteria genomes provided evidence to suggest that these symbionts may play a role in the degradation of host and algal carbohydrates, as well as the storage of phosphate for the host during periods of phosphate limitation(30).

Recently, a third group of ubiquitous sponge-associated betaproteobacterial symbionts were described(42). The proposed new order, the *Tethybacterales*, comprises two families, the *Tethybacteraceae* and the *Persebacteraceae*(42). Based on assessment of MAGs representative of different species within these two families, it was shown that the bacteria within these families had functionally radiated, as they coevolve with their specific sponge host(42). The *Tethybacterales* are distributed both globally and with phylogenetically diverse sponges that represent both high microbial abundance (HMA) and low microbial abundance (LMA) sponges(42). These bacteria have also been detected in other marine invertebrates, ocean water samples, and marine sediment(42) suggesting that these symbionts may have, at one point, been acquired from the surrounding environment.

*Tethybacterales* are conserved in several sponge microbiomes(43) and can be the numerically dominant bacterial population(12, 44–51), with some predicted to be endosymbiotic(12, 44, 49). In *Amphimedeon queenslandica*, the AqS2 symbiont, *Amphirhobacter heronislandensis* (Family Tethybacteraceae), is co-dominant with sulfur-oxidizing *Gammaproteobacteria* AqS1. *A. heronislandensis* AqS2, is present in all stages of the sponge life cycle(52) and appears to have a reduced genome(12). Interestingly, the AqS2 MAG shares some functional similarity with the codominant AqS1, including the potential to generate energy via carbon monoxide oxidation, assimilate sulfur and produce most essential amino acids(12). However, these sympatric symbionts differ significantly in what metabolites they could possibly transport(12).

There are thus two types of sponge-associated symbionts: those that are conserved within a narrow host range, and those that have a broader host range, such as the *SAUL, Entoporibacteria* and *Tethybacterales*. What is currently unknown, is whether these symbionts fulfil the same roles regardless of their host or respond to the needs of the host? In this study we sought to provide answers to these questions, using the dominant, conserved Tethybacterales (strain Sp02-1) symbiont of *Tsitsikamma favus* sponge species (Family *Latrunculiidae*)(53–56) as a springboard into a deeper investigation. Here, we report a comparative study using new and existing *Tethybacterales* genomes and show that functional potential follows that of their taxonomic ranking rather than host-specific adaptation. We also show that the *Tethybacterales* and *Poribacteria* have distinct functional repertoires, that these bacterial families can co-exist in a single host and that the *Tethybacterales* may represent a more ancient lineage of ubiquitous sponge-associated symbionts.

## MATERIALS AND METHODS

### Sponge Collection and taxonomic identification

Sponge specimens were collected by SCUBA or Remotely Operated Vehicle (ROV) from multiple locations within Algoa Bay (Port Elizabeth), the Garden Route National Park, and the Tsitsikamma Marine Protected Area. Collection permits were acquired prior to collections from the Department Environmental Affairs (DEA) and the Department of Environment, Forestry and Fisheries (DEFF) under permit numbers: 2015: RES2015/16 and RES2015/21; 2016:RES2016/11; 2017:RES2017/43; 2018: RES2018/44. Collection metadata are provided in Table S1. Sponge specimens were stored on ice during collection, and thereafter at -20 °C. Subsamples collected for DNA extraction were preserved in RNALater (Invitrogen) and stored at -20 °C. Sponge specimens were dissected, thin sections generated and spicules mounted on microscope slides and examined to allow species identification, as done previously (57–59). Molecular barcoding (28S rRNA gene) was also performed for several of the sponge specimens (Fig. S1) as described previously(53).

### Metagenomic sequencing and analysis

Small sections of each preserved sponge (approx. 2cm^3^) were pulverized in 2ml sterile artificial seawater (24.6 g NaCl, 0.67 g KCl, 1.36 g CaCl_2_.2H_2_O, 6.29 g MgSO_4_.7H_2_O, 4.66 g MgCl_2_.6H_2_O and 0.18 g NaHCO_3_, distilled H_2_O to 1 L) with a sterile mortar and pestle. The resultant homogenate was centrifuged at 16000 rpm for 1 min to pellet cellular material. Genomic DNA (gDNA) was extracted using the ZR Fungal/Bacterial DNA MiniPrep kit (D6005, Zymo Research). Shotgun metagenomic sequencing was performed for four *T. favus* sponge specimens using Ion Torrent platforms. Shotgun metagenomic libraries, of reads 200bp in length, were prepared for each of the four sponge samples respectively (TIC2016-050A, TIC2018-003B, TIC2016-050C and TIC2018-003D) using an Ion P1.1.17 chip. Additional sequence data of 400bp was generated for TIC2016-050A using an Ion S5 530 Chip. TIC2016-050A served as a pilot experiment and we wanted to identify which read length was best for our investigations. However, we did not want to waste additional sequence data and included it when assembling the TIC2016-050A metagenomic contigs so the 400bp reads were included in the assembly of these metagenomes. Metagenomic datasets were assembled into contiguous sequences (contigs) with SPAdes version 3.12.0(60) using the --iontorrent and --only-assembler options. Contigs were clustered into genomic bins using Autometa(61) and manually curated for optimal completion and purity. Validation of the bins was performed using CheckM v1.0.12.(62). Of the 50 recovered genome bins, 5 were of high quality, 13 were of medium quality and 32 were of low quality in accordance with MIMAG standards(63) (Table S2).

### Acquisition and assembly of reference genomes

The genome of *A. queenslandica* symbiont Aqs2 (GCA_001750625.1) was retrieved from the NCBI database. Similarly, other sponge associated *Tethybacterales* MAGs from the JGI database were downloaded and used as references (3300007741_3, 3300007056_3, 3300007046_3, 3300007053_5, 3300021544_3, 3300021545_3, 3300021549_5, 2784132075, 2784132054, 2814122900, 2784132034 and 2784132053).

Thirty-six raw read SRA datasets from sponge metagenomes were downloaded from the SRA database (Table S3). Illumina reads from these datasets were trimmed using Trimmomatic v0.39(64) and assembled using SPAdes v3.14(60) in --meta mode and resultant contigs were binned using Autometa(61). This resulted in a total of 393 additional genome bins, the quality of which was assessed using CheckM(62) and taxonomically classified with GTDB-Tk(65) with database release95 (Table S2). A total of 27 bins were classified as AqS2 and were considered likely members of the newly proposed *Tethybacterales* order(42). However, 10 of the 27 bins were low quality and were not used in downstream analyses. In addition, 59 *Poribacteria* genome bins were downloaded from the NCBI database for functional comparison (Table S4) and three were used from the 393 genome bins generated in this study (*Geodia parva* sponge hosts).

### Taxonomic identification

Partial and full-length 16S rRNA gene sequences were extracted from bins using barrnap 0.9 (https://github.com/tseemann/barrnap). Extracted sequences were aligned against the nr database using BLASTn(66). Genomes were additionally uploaded individually to autoMLST(67) and analyzed in both placement mode and *de novo* mode (IQ tree and ModelFinder options enabled, and concatenated gene tree selected). All bins and downloaded genomes were taxonomically identified using GTDB-Tk(65).

### Genome annotation and metabolic potential analysis

All bins and downloaded genomes were annotated using Prokka 1.13(68) with NCBI compliance enabled. Protein-coding amino-acid sequences from genomic bins were annotated against the KEGG database using kofamscan(69) with output in mapper format. Custom Python scripts were used to summarize annotation counts (Find scripts here: https://github.com/samche42/Family_matters). Potential Biosynthetic Gene Clusters (BGCs) were identified by uploading genome bins to the antiSMASH web server(70) with all options enabled. Predicted amino acid sequences of genes within each identified gene cluster were aligned against the nr database using BLASTn(66) to identify the closest homologs. Protein sequences of genes within each identified gene cluster were aligned against the nr database using BLASTn(66) to identify the closest homolog.

### Phylogeny and function of *Tethybacterales* species

A subset of orthologous genes common to all medium quality *Tethybacterales* genomes/bins was created. Shared amino acid identity (AAI) was calculated with the aai.rb script from the enveomics package(71). 16S rRNA genes were analyzed using BLASTn(66). Functional genes were annotated against the KEGG database using kofamscan(69). Annotations were collected into functional categories and visualized in R (See https://github.com/samche42/Family_matters for all scripts). A Non-metric Multidimensional Scaling (NMDS) plot of the presence/absence metabolic counts was constructed using Bray-Curtis distance using the vegan package(72) in R. Analysis of Similarity (ANOSIM) analyses were also conducted using the vegan package in R using Bray-Curtis distance and 9999 permutations.

### Genome divergence estimates

Divergence estimates were performed as described previously(73). Briefly, homologous genes in *Tethybacterales* genomes were identified using OMA v. 2.4.2.(74). A subset of homologous genes present in all genomes was created. Homologous genes were aligned using MUSCLE v.3.8.155(75) and clustered into fasta files representing each genome using merge_fastas_for_dNdS.py (See https://github.com/samche42/Family_matters for all scripts). Corresponding nucleotide sequences extracted from PROKKA annotations using multifasta_seqretriever.py. All stop codons were removed using remove_stop_codons.py. All nucleotide sequences, per genome, were concatenated to produce a single nucleotide sequence per genome using the union function from EMBOSS(76).

All amino acid sequences were similarly concatenated. This resulted in a single concatenated nucleotide sequence and a single concatenated amino acid sequence per genome. Concatenated nucleotide sequences were clustered into two fasta files (one nucleotide, one protein sequence) and then aligned using PAL2NAL(77). The resultant alignment was then run in codeml to produce pairwise synonymous substitution rates (dS). Divergence estimates can be determined by dividing pairwise dS values by a given substitution rate, and further divided by 1 million. Pairwise synonymous substitution rates can be found in Table S5. Pairwise divergence values were illustrated as a tree using MEGAX(78). Concatenated amino acid and nucleotide sequences of the 18 orthologous genes were aligned using MUSCLE v.3.8.155(75) and the evolutionary history inferred using the UPMGA method(79) in MEGAX(78) with 10000 bootstrap replicates.

### Identification of unique and host-associated genes in putative symbiont genome bins

A custom database of genes from all bacterial bins (with the exception of the *Tethybacterales* Sp02-1 symbionts) was created using the “makedb” option in Diamond(80) to identify genes that were unique to the Sp02-1 symbionts. To be exhaustive and screen against the entire metagenome, genes from low-quality genomes (except low-quality Sp02-1 genomes), small contigs (<3000 bp) that were not included in binning and unclustered contigs (i.e., included in binning but were not placed within a bin) were included in this database. Sp02-1 genes were aligned using diamond blast(80). A gene was considered “unique” if the aligned hit shared less than 40% amino acid identity with any other genes from the *T. favus* metagenomes and had no significant hits against the nr database or were identified as pseudogenes. All “unique” Sp02-1 genes annotated as “hypothetical” (both Prokka and NCBI nr database annotations) were removed. Finally, we compared Prokka annotation strings between the three Sp02-1 genes and all other *T. favus* associated genome bins and excluded any Sp02-1 genes that were found to have the same annotation as a gene in one of the other bins.

## DATA AVAILABILITY

All data can be accessed from the NCBI website under BioProject PRJNA508092.

## RESULTS AND DISCUSSION

The microbiomes of sponges of the *Latrunculiidae* family are highly conserved and are dominated by populations of related betaproteobacterial symbionts. Based on their 16S rRNA gene sequence, the *Betaproteobacteria* Sp02-1 symbionts from different latrunculid sponges are closely related and are likely members of the newly described *Tethybacterales*.

### Characterization of putative betaproteobacteria Sp02-1 genome bin

Genome bin 003B_4, from sponge specimen TIC2018-003B, included a 16S rRNA gene sequence that shared 99.86% identity with the *T. favus*-associated *Tethybacterales* Sp02-1 full-length 16S rRNA gene clone (HQ241787.1). The next closest relatives were uncultured betaproteobacterium 16S clones from a *Xestospongia muta* and *Tethya aurantium* sponges (Fig. S2). Bins 050A_14, 050C_6 and 003D_6 were also identified as possible representatives of the *Tethybacterales* Sp02-1 based on their predicted phylogenetic relatedness. However, they were of low quality and not used in downstream analyses.

Genome bin 003B_4 was used as a representative of the *Tethybacterales* Sp02-1 symbiont. Bin 003B_4 is approximately 2.95 Mbp in size and of medium quality per MIMAG standards(63) (Table S2), and it has a notable abundance of pseudogenes (∼25% of all genes), which resulted in a coding density of 65.27%, far lower than the average for bacteria(81). An abundance of pseudogenes and low coding density, is usually an indication that the genome in question may be undergoing genome reduction (82) similar to other betaproteobacteria in the proposed order of *Tethybacterales*(42).

The *Tethybacterales* Sp02-1 genome encodes all genes necessary for glycolysis, PRPP biosynthesis and most genes required for the citrate cycle and oxidative phosphorylation were detected in the gene annotations. Also present are the genes necessary to biosynthesize valine, leucine, isoleucine, tryptophan, phenylalanine, tyrosine and ornithine amino acids as well as genes required for transport of L-amino acids, proline and branched amino acids. This would suggest that this bacterium may exchange amino acids with the host, as observed previously in both insect and sponge-associated symbioses(8, 83, 84).

A total of 13 genes unique to the *Tethybacterales* Sp02-1 symbiont were identified (i.e., not identified elsewhere in the *T. favus* metagenomes). One gene was predicted to encode an ABC transporter permease subunit that was likely involved in glycine betaine and proline betaine uptake. A second gene encoded 5-oxoprolinase subunit PxpA (Table S6). The presence of these two genes suggests that the *Tethybacterales* Sp02-1 genome can acquire proline and convert it to glutamate(85) in addition to glutamate already produced via glutamate synthase. Other unique genes encode a restriction endonuclease subunit and site-specific DNA-methyltransferase, which would presumably aid in defense against foreign DNA. At least seven of the unique gene products are predicted to be associated with phages, including the anti-restriction protein ArdA. ArdA is a protein that has previously been shown to mimic the structures of DNA normally bound by Type I restriction modification enzymes, which prevent DNA cleavage, and effectively results in anti-restriction activity(86). If functionally active in the *Tethybacterales* Sp02-1 symbiont, we speculate that this protein may similarly prevent DNA cleavage through its mimicry of the targeted DNA structures and protect the genome against Type I restriction modification enzymes. Finally, two of the unique genes were predicted to encode an ankyrin repeat domain-containing protein and a VWA domain-containing protein. These two proteins are known to be involved in cell-adhesion and protein-protein interactions(87, 88) and if active within the symbiont, they may help facilitate the symbiosis between the *Tethybacterales* Sp02-1 symbiont and the sponge host.

### Comparison of putative Sp02-1 with other Tethybacterales

Several betaproteobacteria (*Tethybacterales*) sponge symbionts have been described to date and these bacteria are thought to have functionally diversified following the initiation of their ancient partnership(42). To test this hypothesis, we downloaded twelve genomes/MAGs of *Tethybacterales* (classified as AqS2 in GTDB) from the JGI database. Additionally, we assembled and binned metagenomic data from thirty-six sponge SRA datasets, covering fourteen sponge species and recovered an additional fourteen AqS2-like genomes. Ten of the total twenty-seven bins were of low quality, so Bin 003B_4 (Sp02-1) and sixteen medium quality *Tethybacterales* bins/genomes were used for further analysis (Table 1).

**Table 1.** General characteristics of putative Tethybacterales genomes/MAGs.

| Genome | Sponge host | Size (Mbp) | Complete (%) | Contam (%) | Quality | “Core” genes | Study/ Accession |
| --- | --- | --- | --- | --- | --- | --- | --- |
| “*Candidatus* Ukwabelana africanus” (003B_4) | *Tsitsikamma favus* | 2.96 | 72.92 | 3.56 | Medium | 84.52 | This study |
| "*Candidatus* Regalo mexicanus" (ImetM1_9) | *Iophon methanophila* | 1.56 | 85.58 | 0.61 | Medium | 83.33 | This study |
| "*Candidatus* Regalo mexicanus" (ImetM2_1_1) | *Iophon methanophila* | 1.6 | 84.36 | 0.61 | Medium | 86.9 | This study |
| *Perseobacter sydneyensis* (C29) | *Crella incrustans* | 1.52 | 82.87 | 0.61 | Medium | 78.57 | Taylor et al., 2021<br>JGI: 2784132075 |
| *Tethyobacter castellensis* (ccb2r) | *Cymbastela concentrica* | 1.69 | 81 | 0.3 | Medium | 85.71 | Taylor et al., 2021<br>JGI: 2784132054 |
| *Beroebacter blansensis* (Crambe1) | *Crambe crambe* | 2.25 | 80.38 | 1.81 | Medium | 73.8 | Taylor et al., 2021<br>JGI: 2814122900 |
| *Amphirhobacter heronislandensis* (AqS2) | *Amphimedon queenslandica* | 1.61 | 71.13 | 1.52 | Medium | 80.95 | Gauthier et al., 2016<br>NCBI: GCA_001750625.1 |
| *Calypsobacter congwoongensis* (B3) | *Scopalina* sp. | 1.05 | 64.33 | 0.04 | Medium | 53.57 | Taylor et al., 2021<br>JGI: 2784132034 |
| *Telestobacter tawharauni* (TSB1) | *Tethya stolonifera* | 1.24 | 56.3 | 0 | Medium | 57.14 | Taylor et al., 2021<br>JGI: 2784132053 |
| "*Candidatus* Hadiah malacca" (Csing_1) | *Coelocarteria singaporensis* | 1.58 | 65.47 | 0.61 | Medium | 73.81 | JGI: 3300007741_3 |
| "*Candidatus* Hadiah malacca" (Csing_2) | *Coelocarteria singaporensis* | 1.36 | 58.12 | 0 | Medium | 72.62 | JGI: 3300007056_3 |
| "*Candidatus* Hadiah malacca" (Csing_3) | *Coelocarteria singaporensis* | 1.44 | 57.04 | 0.61 | Medium | 76.19 | JGI: 3300007046_3 |
| "*Candidatus* Hadiah malacca" (Csing_5) | *Coelocarteria singaporensis* | 1.15 | 55.82 | 0.61 | Medium | 75.00 | JGI: 3300021544_3 |
| "*Candidatus* Hadiah malacca" (Csing_6) | *Coelocarteria singaporensis* | 1.36 | 58.12 | 0 | Medium | 72.62 | JGI: 3300021545_3 |
| "*Candidatus* Hadiah malacca" (Csing_7) | *Coelocarteria singaporensis* | 0.98 | 54.76 | 1.93 | Medium | 69.05 | JGI: 3300021549_5 |
| "*Candidatus* Dora taiwanensis" (CCyA_2_3) | *Cinachyrella* sp. | 3.08 | 84.92 | 1.83 | Medium | 78.57 | This study |
| "*Candidatus* Dora taiwanensis" (CCyB_3_2) | *Cinachyrella* sp. | 3.45 | 79.58 | 5.93 | Medium | 75.00 | This study |
| CCyA_16_0 | *Cinachyrella* sp. | 0.86 | 16.95 | 3.81 | Low | 35.71 | This study |
| CCyA_3_0 | *Cinachyrella sp.* | 0.62 | 29.31 | 0 | Low | 15.48 | This study |
| CCyA_3_39 | *Cinachyrella sp.* | 0.41 | 22.01 | 4.88 | Low | 36.90 | This study |
| CCyB_501_11 | *Cinachyrella sp.* | 0.04 | 7.51 | 0 | Low | 17.86 | This study |
| CCyB_502_83 | *Cinachyrella sp.* | 0.43 | 17.01 | 0 | Low | 21.43 | This study |
| CCyC_2_7 | *Cinachyrella sp.* | 1.81 | 47.5 | 0 | Low | 57.14 | This study |
| 050C_6 | *Tsitsikamma favus* | 2.21 | 24.03 | 4.51 | Low | 32.14 | This study |
| Csing_4 | *Coelocarteria singaporensis* | 1.07 | 39.35 | 0.61 | Low | 54.76 | JGI: 3300007053_5 |
| 050A_14 | *Tsitsikamma favus* | 6.08 | 61 | 19.4 | Low | 73.81 | This study |
| 003D_6 | *Tsitsikamma favus* | 0.32 | 0 | 0 | Low | 0 | This study |

First, the phylogeny of the *Tethybacterales* symbionts was determined using single-copy marker genes in autoMLST, revealing a deep branching clade of these sponge-associated symbionts and that bin 003B_4 clustered within the proposed *Persebacteraceae* family (Fig. 1). All members of the *Persebacteraceae* family dominate the microbial community of their respective sponge hosts(44, 53, 54, 89). We additionally identified what appears to be a third family, consisting of symbionts associated with *C. singaporensis* and *Cinachyrella* sponge species (Fig. 1). Assessment of shared AAI (Average Amino acid Identity) indicates that these genomes represent a new family, sharing an average of 80% AAI within the family (Table S7)(90). These three families share less than 89% sequence similarity with respect to their 16S rRNA sequences, with intra-clade differences of less than 92% (Table S8). Therefore, they may represent novel classes within the *Tethybacterales* order (90). In keeping with naming the families after Oceanids of Greek mythology(42), we propose the family name *Polydorabacteraceae*, which means “many gifts”. Additionally, we propose species names for the newly identified genera as follows: Bin 003B_4 is a single representative of “*Candidatus* Ukwabelana africanus”, Bin Imet_M1_9 and Bin ImetM2_1_1 are both representatives of “*Candidatus* Regalo mexicanus”, Bin CCyA_2_3 and CCyB_3_2 are both representatives of “*Candidatus* Dora taiwanensis”, and all six bins from *C. singaporensis* are representative of “*Candidatus* Hadiah malacca”. In each case, the genus name means “gift from” in the local language (where possible) from where the host sponge was collected, and the species name reflects the region/country from which the sponge host was collected.

**Figure 1.**
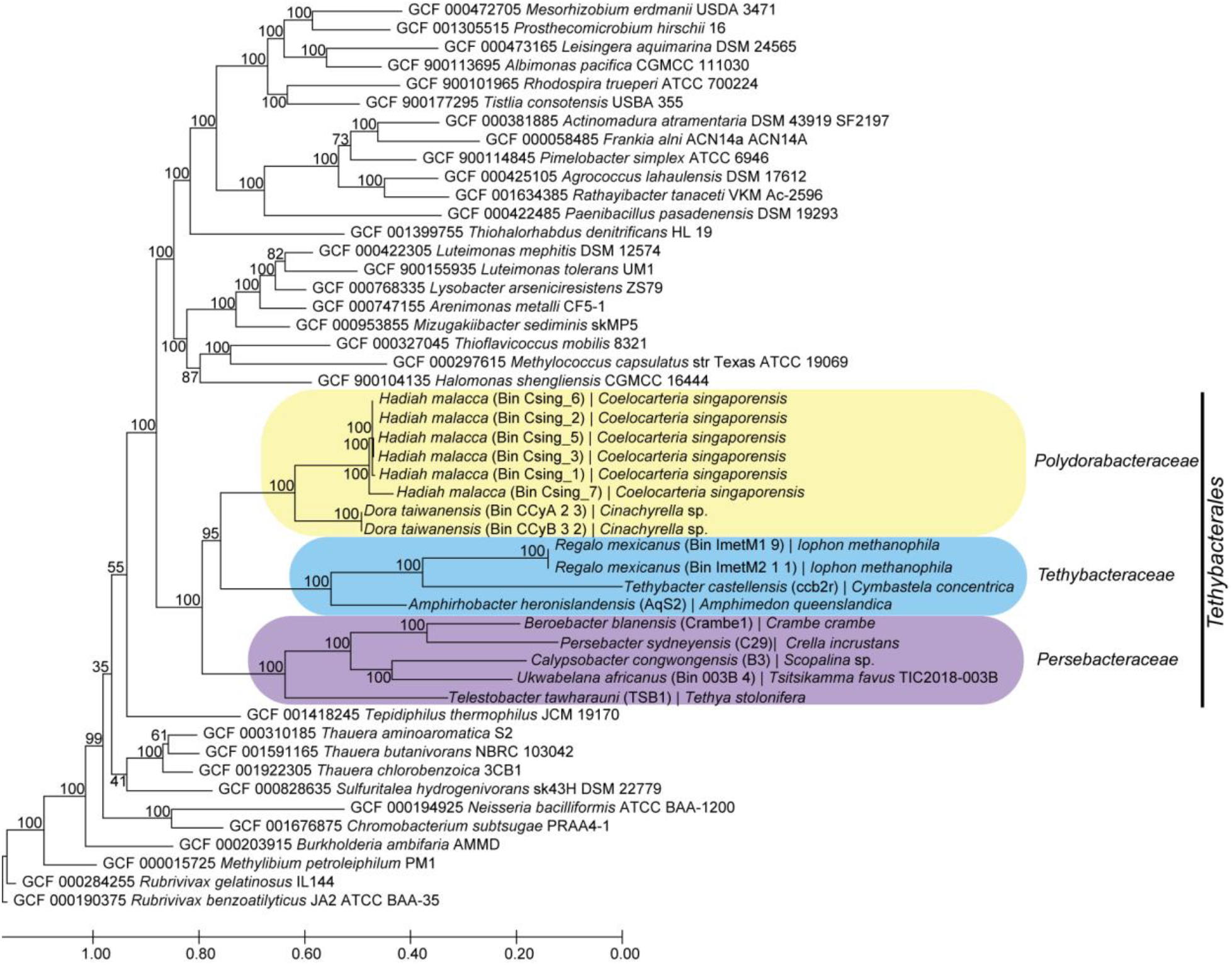
Phylogeny of the *Tethybacterales* sponge symbionts. Using autoMLST, single-copy markers were selected and used to delineate the phylogeny of these sponge-associated betaproteobacteria revealing a new family of symbionts in the *Tethybacterales* order. Additionally, it was shown that the *T. favus* associated Sp02-1 symbiont belongs to the *Persebacteraceae* family. The phylogenetic tree was inferred using the de novo method in AutoMLST using a concatenated alignment with IQ Tree and ModelFinder enabled. Branch lengths are proportional to the number of substitutions per site.

We identified 4306 groups of orthologous genes between all seventeen *Tethybacterales* genomes, with only eighteen genes common to all the genomes. More shared genes were expected, but as several of the genomes investigated are incomplete, it is possible that additional common genes would be found if the genomes were complete. Hierarchical clustering of gene presence/absence data revealed that the gene pattern of Bin 003B_4 most closely resembled that of *Tethybacterales* genomes from *C. crambe, C. incrustans* and the *Scopalina* sp. sponges (family *Persebacteraceae*) (Fig. 2A). Thirteen of the shared genes between all *Tethybacterales* genomes encoded ribosomal proteins or those involved in energy production. Genes encoding chorismate synthase were found across all seventeen genomes and suggest that tryptophan production may be shared among these bacteria. According to a recent study, *Dysidea etheria* and *A. queenslandica* sponges cannot produce tryptophan (a possible essential amino acid), which may indicate a common role for the *Tethybacterales* symbionts as tryptophan producers(91). Several other shared genes were predicted to encode proteins involved in stress responses, including protein-methionine-sulfoxide reductase, ATP-dependent Clp protease and chaperonin enzyme proteins, which aid in protein folding or degradation under various stressors(92–96). Internal changes in oxygen levels(97), and temperature changes(98–100) are examples of stressors experienced by the sponge holobiont. It is unsurprising that this clade of largely sponge-specific *Tethybacterales* share the ability to deal with these many stressors as they adapt to their fluctuating environment.

**Figure 2.**
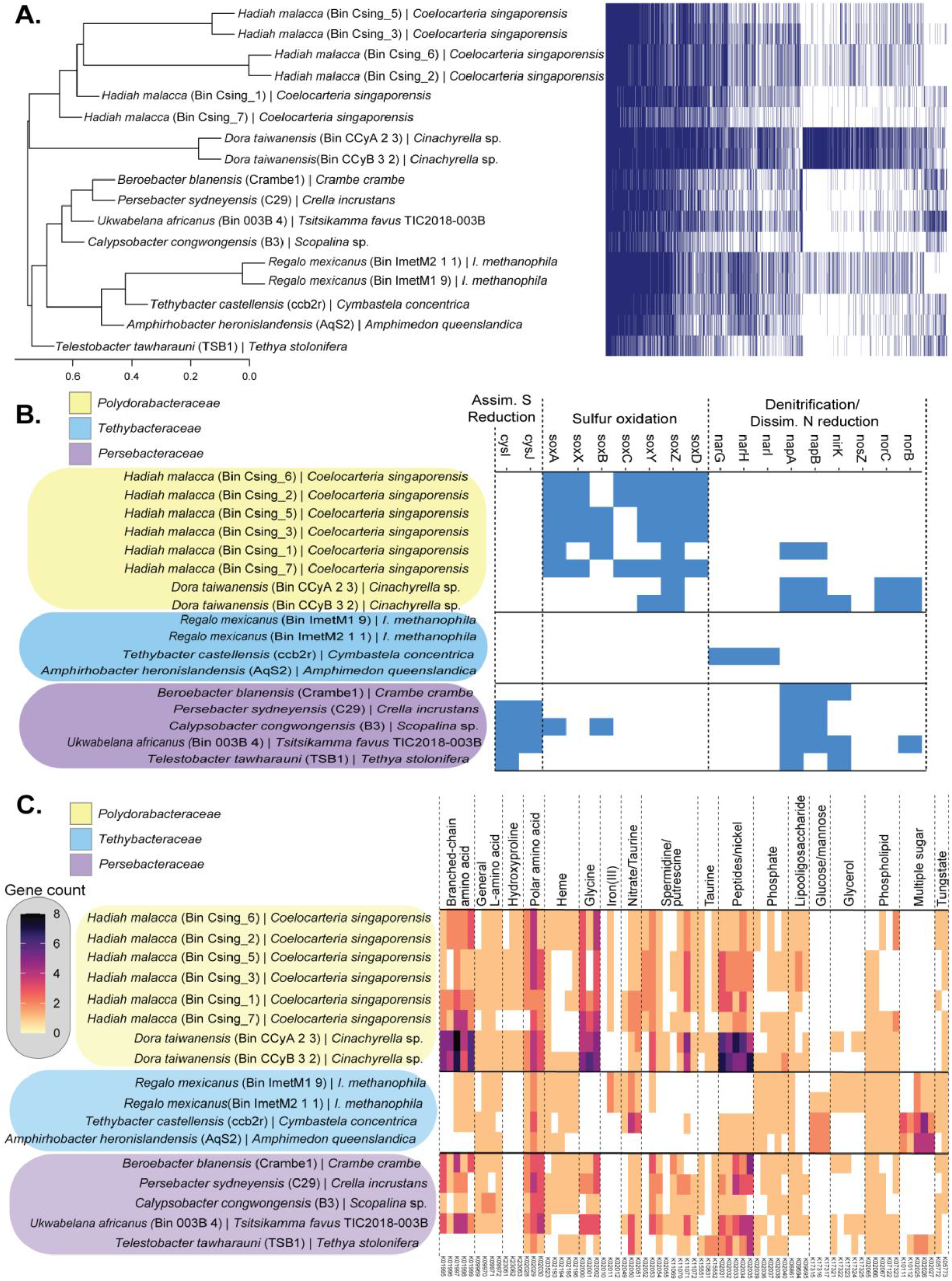
Functional specialization of *Tethybacterales* families. The newly proposed *Tethybacterales* order appears to consist of three bacterial families. These families appear to have similar gene distribution (A) where the potential function of these genes indicates specialization in nutrient cycling (B) and solute transport (C).

Alignment against the KEGG database revealed some noteworthy trends that differentiated the three *Tethybacterales* families (Fig. 2B, Table S9): (1) the genomes of the proposed *Polydorabacteraceae* family include several genes associated with sulfur oxidation; (2) the *Persebacteraceae* are unique in their potential for reduction of sulfite (*cysIJ*) and (3) the *Tethybacteraceae* have the potential for cytoplasmic nitrate reduction (*narGHI*), while the other two families may perform denitrification. Similarly, the families differ to some extent in what can be transported in and out of the symbiont cell (Fig. 2C). Proposed members of the *Polydorabacteraceae* appear exclusively capable of transporting hydroxyproline which may imply a role in collagen degradation(101). The *Tethybacteraceae* and *Persebacteraceae* appear able to transport spermidine, putrescine, taurine, and glycine which, in combination with their potential to reduce nitrates, may suggest a role in C-N cycling(102). All three families transport various amino acids as well as phospholipids and heme. The exchange of amino acids between symbiont and sponge host has previously been observed(103) and may provide the *Tethybacterales* with a competitive advantage over other sympatric microorganisms(104) and possibly allow the sponge hosts to regulate the symbioses via regulation of the quantity of amino acids available for symbiont uptake(105). Similarly, the transfer of heme in the iron-starved ocean environment between sponge host and symbiont could provide a selective advantage as heme may act as a supply of iron(106). The *Tethybacteraceae* were distinct from the other two families in their potential to transport sugars. As mentioned earlier, the transport of sugars plays an important role in symbiotic interactions(98, 107–109) and it is possible that this family of symbionts require sugars from their sponge hosts.

### Comparative analyses of functional potential between *Tethybacterales* and *Poribacteria*

We wanted to determine whether broad-host range sponge-associated symbionts have converged to perform similar roles in their sponge hosts. Accordingly, we annotated 62 *Poribacteria* genomes which consisted of 24 *Pelagiporibacteria* (free-living) and 38 *Entoporibacteria* (sponge-associated) genomes, and the 17 *Tethybacterales* genomes against the KEGG database. We catalogued the presence/absence of 896 unique genes spanning carbohydrate metabolism, methane metabolism, nitrogen metabolism, sulfur metabolism, phosphate metabolism and several transporter systems (Fig. 3, Table S9). Inspection of the functional potential in the *Tethybacterales* and *Poribacteria* revealed several insights (Fig. 3). The gene repertoires of the *Poribacteria* and the *Tethybacterales* are distinct from one another (Fig. S3, Table 2), with notable differences including the genes associated with denitrification, dissimilatory nitrate reduction, thiosulfate oxidation, hydroxyproline transport, glycine betaine/proline transport, glycerol transport, taurine transport, tungstate transport and glucose/mannose transport, all of which are present in the *Tethybacterales* and absent in the *Poribacteria* (Fig. 3). Conversely, several gene clusters were detected in the *Poribacteria* and absent in the *Tethybacterales*, including trehalose biosynthesis, the Entner-Doudoroff pathway, galactose degradation, phosphate metabolism, phosphonate transport, assimilatory sulfate reduction, molybdate transport, osmoprotectant transport, hydroxymethylpyrimidine transport (Fig. 3). It has been reported that both *Entoporibacteria* and *Pelagiporibacteria* include genes associated with denitrification(33), however we could not detect many genes associated with nitrogen metabolism in our analyses (Fig. 3).

**Figure 3.**
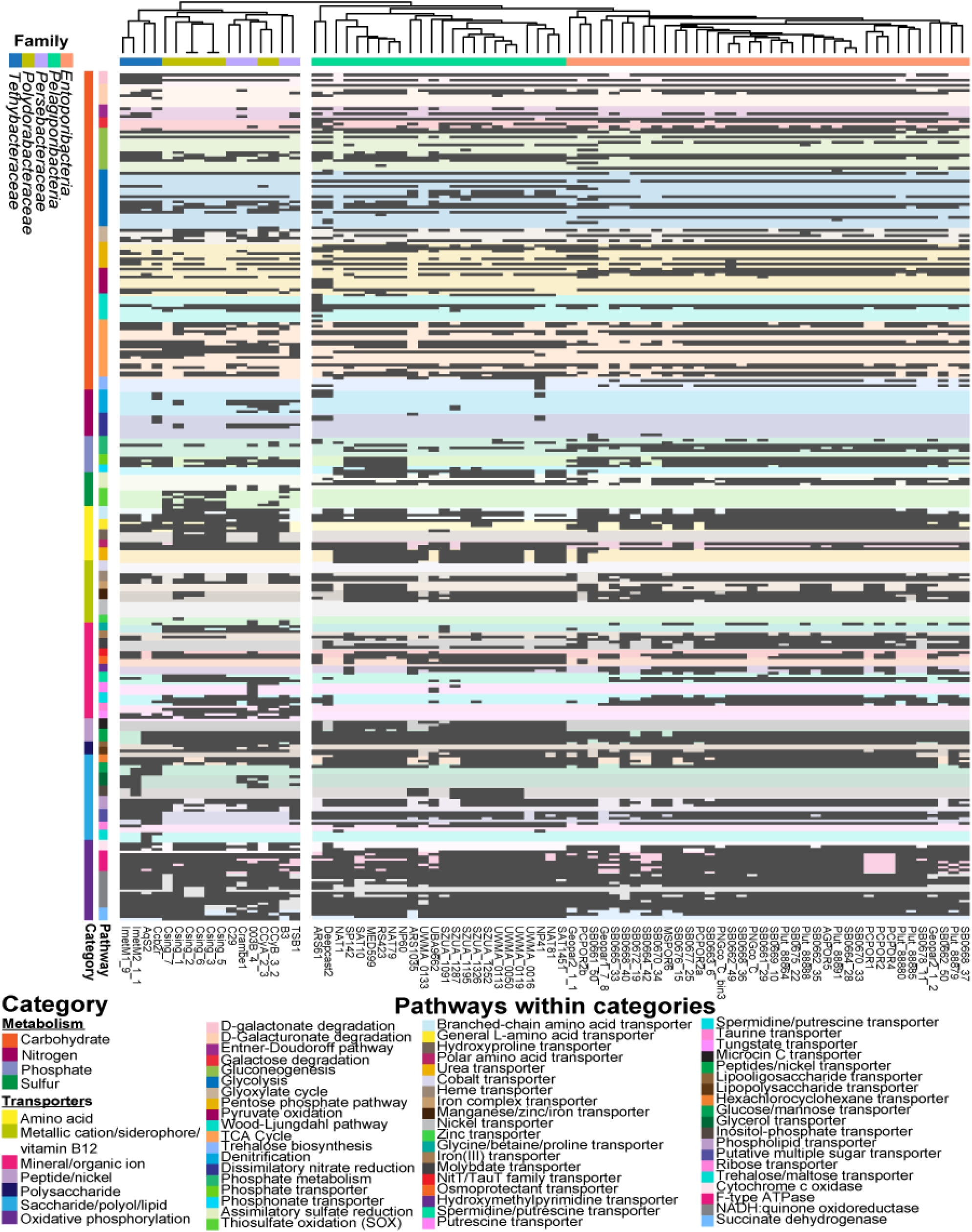
Functional differences between *Tethybacterales* and *Poribacteria*. Sponge-associated *Tethybacterales* genomes include significantly different functional gene repertoires to those found in *Poribacteria*. Detailed presence/absence of metabolic genes (KEGG annotations) detected in *Tethybacterales* and *Poribacteria* genome bins. Genomes are listed at the bottom of the figure and clustered according to the presence/absence of functional genes (top). Taxonomic classification of each genome is indicated using a colored bar (top). Functional genes are collected into their respective pathways, which are further organized into larger functional categories. A colored key is provided (right) and is in the same order as that of the colored blocks in the graph (left).

**Table 2:** Pairwise ANOSIM of presence/absence of KEGG-annotated functional genes in Poribacteria and Tethybacterales.

| Taxon A | Taxon B | p-value | R statistic |
| --- | --- | --- | --- |
| *Entoporiobacteria* | *Polydorabacteraceae* | 0.0001 | 0.9965 |
| *Entoporiobacteria* | *Pelagiporiobacteria* | 0.0001 | 0.8881 |
| *Entoporiobacteria* | *Perseobacteraceae* | 0.0001 | 0.9985 |
| *Entoporiobacteria* | *Tethybacteraceae* | 0.0001 | 0.9938 |
| *Polydorabacteraceae* | *Pelagiporiobacteria* | 0.0001 | 0.9951 |
| *Polydorabacteraceae* | *Perseobacteraceae* | 0.0045 | 0.3974 |
| *Polydorabacteraceae* | *Tethybacteraceae* | 0.0025 | 0.9908 |
| *Pelagiporiobacteria* | *Perseobacteraceae* | 0.0001 | 0.9961 |
| *Pelagiporiobacteria* | *Tethybacteraceae* | 0.001 | 0.9741 |
| *Perseobacteraceae* | *Tethybacteraceae* | 0.0085 | 0.7500 |

We cross-checked gene annotations generated using Prokka (HAMAP database) and Blast (nr database). Genes associated with assimilatory nitrate reduction (*narB*, and *nirA*) were identified in *Poribacteria* using these alternate annotations, but we could not detect genes associated with denitrification in the *Poribacteria*. Conversely, genes associated with denitrification (*napAB* and *nirK*) were detected in the *Persebacteraceae* of the Tethybacterales in Prokka, Blast and KEGG annotations (Fig. 3), indicating that their absence in *Poribacteria* genomes was not an artefact of our analyses.

Pairwise ANOSIM analysis (using Bray-Curtis distance) confirmed that the functional genetic repertoire (KEGG annotations) of the *Tethybacterales* bacteria showed a strong, significant dissimilarity to that of the sponge-associated *Entoporibacteria* and the free-living *Pelagiporibacteria* (Table 2). In addition, the *Polydorabacteraceae* and the *Persebacteraceae* were significantly different from one another, but the lower R-statistic would suggest that the dissimilarity is not as strong as between other groups in this analysis, while the *Tethybacteraceae* appear to be more functionally distinct from the other two *Tethybacterales* families.

Taken together, these data suggest that the three *Tethybacterales* families and the *Entoporibacteria* lineages may each fulfil distinct functional or ecological niches within a given sponge host. Interestingly, some sponges such as *A. aerophoba* and *C. singaporenesis*, can play host to both *Tethybacterales* and *Entoporibacteria* species (Table S2) which provides further evidence that these symbionts may serve different purposes within their sponge host.

We investigated the respective approximate divergence pattern of the *Tethybacterales* and the *Entoporibacteria* and whether their divergence followed that of their sponge hosts. The eighteen homologous genes shared between the *Tethybacterales* were used to estimate the rate of synonymous substitution, which provides an approximation for the pattern of divergence between the species(110). We found that the estimated divergence pattern of the Tethybacterales (Fig. 4A) and the phylogeny of the host sponges (Fig. 4B) was incongruent. Phylogenetic trees inferred using single-copy marker genes (Fig. 1), and the comprehensive 16S rRNA tree published by Taylor and colleagues(42) confirm this lack of congruency between symbiont and host phylogeny. Other factors such as collection site or depth could not explain the observed trend. Similar incongruence of symbiont and host phylogeny was observed for the *Entoporibacteria* (34 homologous genes used to estimate synonymous substitution rates) (Fig. 4C-D), in agreement with previous phylogenetic studies (31, 33, 34). This would suggest that these sponges likely acquired a free-living *Tethybacterales* common ancestor at different time points throughout their evolution, and that the same is true for the *Entoporibacteria*. Evidence of coevolution of betaproteobacteria symbionts within sponges families (46, 52, 53, 111) implies that *Tethybacterales* symbionts were likely acquired horizontally at various time points and may have coevolved with their respective hosts subsequent to acquisition.

**Figure 4.**
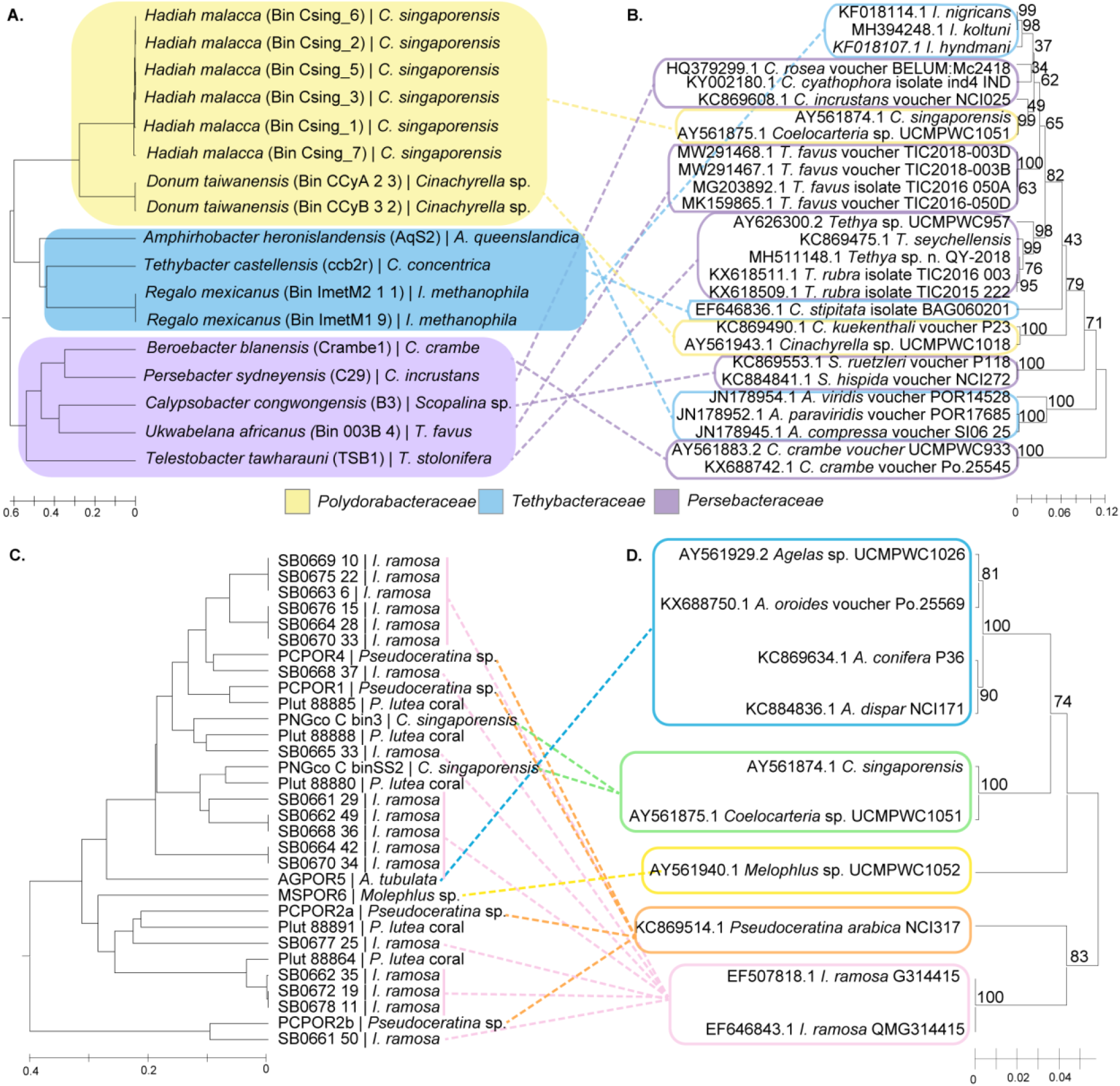
The divergence pattern of sponge-associated *Tethybacterales, Entoporibacteria* and their respective host sponges. The divergence of the *Tethybacterales* and *Entoporibacteria* is incongruent with the phylogeny of the host sponges. (A and C) Branch length of symbiont divergence estimates is proportional to the pairwise rate of synonymous substitution calculated (ML estimation) using a concatenation of genes common to all genomes. Rate of synonymous substitution was calculated using PAL2NAL and CodeML from the PAML package and visualized in MEGAX. (B and D) Phylogeny of host sponges (or close relatives thereof) was inferred with 28S rRNA sequence data using the UPGMA method and Maximum Composite Likelihood model with 1000 bootstrap replicates. Branch lengths indicate the number of substitutions per site. All ambiguous positions were removed for each sequence pair (pairwise deletion option). Evolutionary analyses were conducted in MEGA X.

Finally, the estimated rates of synonymous substitution of homologous genes were used to estimate the relative times at which the *Tethybacterales* and *Entoporibacteria* taxa began diverging, respectively. Regardless of substitution rate used, it was found that the sponge-associated *Tethybacterales* genomes began diverging from one another before the *Entoporibacteria* began diverging from one another (Table S5). If one accepts that divergence between exclusively sponge-associated bacterial lineages began when the common ancestor first associated with a sponge host, then the earlier divergence of sponge-associated *Tethybacterales* (relative to the *Entoporibacteria*) suggests that the *Tethybacterales* may have associated with sponges before the Poribacteria common ancestor and represent a more ancient symbiont. However, this hypothesis may prove false if additional *Entoporibacteria* lineages are discovered and added to the analyses, or other factors such as mutation rates, time between symbiont acquisition and transition to vertical inheritance of symbionts or fossil records disprove this hypothesis.

## CONCLUSION

Here, we have shown that the family to which a broad host-range symbiont belongs, such as the *Tethybacterales*, dictates the functional potential of the symbiont, whereas in narrow host-range symbionts, potential function is often a strain-specific trait. This work has expanded our understanding of the *Tethybacterales* and the possible functional specialization of the families within this new order. The *Tethybacterales* are functionally distinct from the *Poribacteria* which would suggest that although these bacteria are both ubiquitously associated with a wide range of sponge hosts, they likely have not converged to fulfil the same role. Instead, it would appear that these symbionts were selected by the various sponge hosts for existing functional capabilities. The incongruence of both *Tethybacterales* and *Entoporibacteria* suggests that their ancestors were horizontally acquired at different evolutionary timepoints, and co-evolution may have occurred following the establishment of the association. Estimates of when the *Tethybacterales* and *Entoporibacteria* began diverging from their respective common ancestors implied that *Tethybacterales* may have associated with a sponge host before the *Entoporibacteria* and therefore the Tethybacterales may be an older sponge-associated symbiont. However, additional data is required to validate or disprove this hypothesis.

## ACKNOWLEDGEMENTS

This research was performed in part using the computer resources and assistance of the UW-Madison Center for High Throughput Computing (CHTC) in the Department of Computer Sciences. The CHTC is supported by UW-Madison, the Advanced Computing Initiative, the Wisconsin Alumni Research Foundation, Wisconsin Institutes for Discovery and the National Science Foundation and is an active member of the Open Science Grid, which is supported by the National Science Foundation and the U.S. Department of Energy’s Office of Science. The authors also acknowledge the Center for High-Performance Computing (CHPC, South Africa) for providing computing facilities for bioinformatics data analysis. The authors acknowledge Gwynneth Matcher, Mr. Carel van Heerden and Ms. Alvera Vorster for their NGS technical support. The authors thank Ryan Palmer, Skipper Koos Smith, Nicholas Riddin and Nicholas Schmidt (ACEP) for technical support and expertise during sponge collections. We thank the South African Environmental Observation Network (SAEON), Elwandle Coastal Node, and the Shallow Marine and Coastal Research Infrastructure (SMCRI) for the use of their research platforms and infrastructure and South African National Parks (SANParks) for their assistance and support.

## SUPPLEMENTARY FIGURES

**Figure S1.**
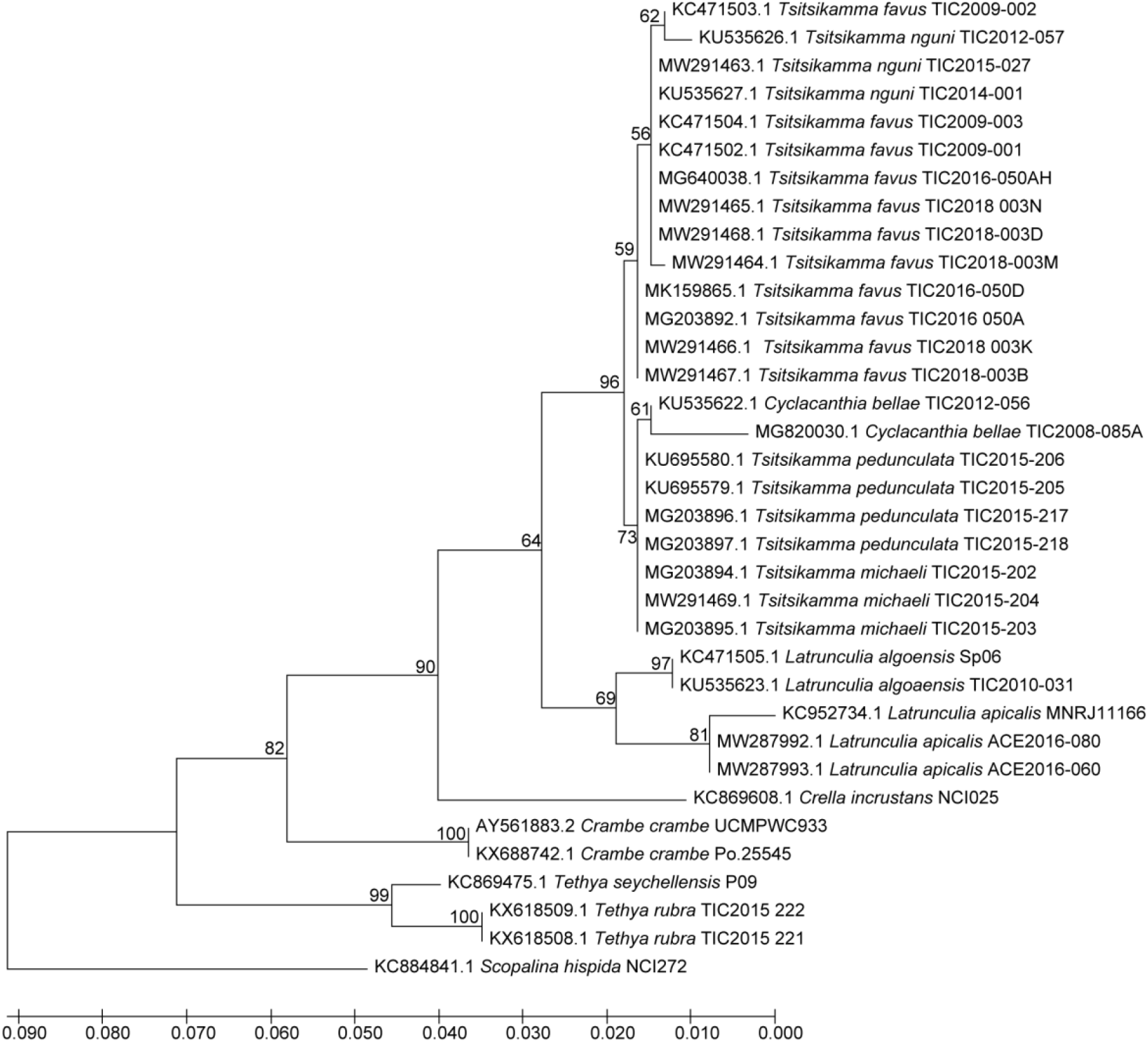
Phylogeny of sponges within the family *Latrunculiidae*. Sponge phylogeny was inferred using 28S rRNA sequence data using the Maximum-likelihood method and Tamura-Nei model with 1000 bootstrap replicates. Branch lengths indicate the number of substitutions per site. All ambiguous positions were removed for each sequence pair (pairwise deletion option). Evolutionary analyses were conducted in MEGA X.

**Figure S2.**
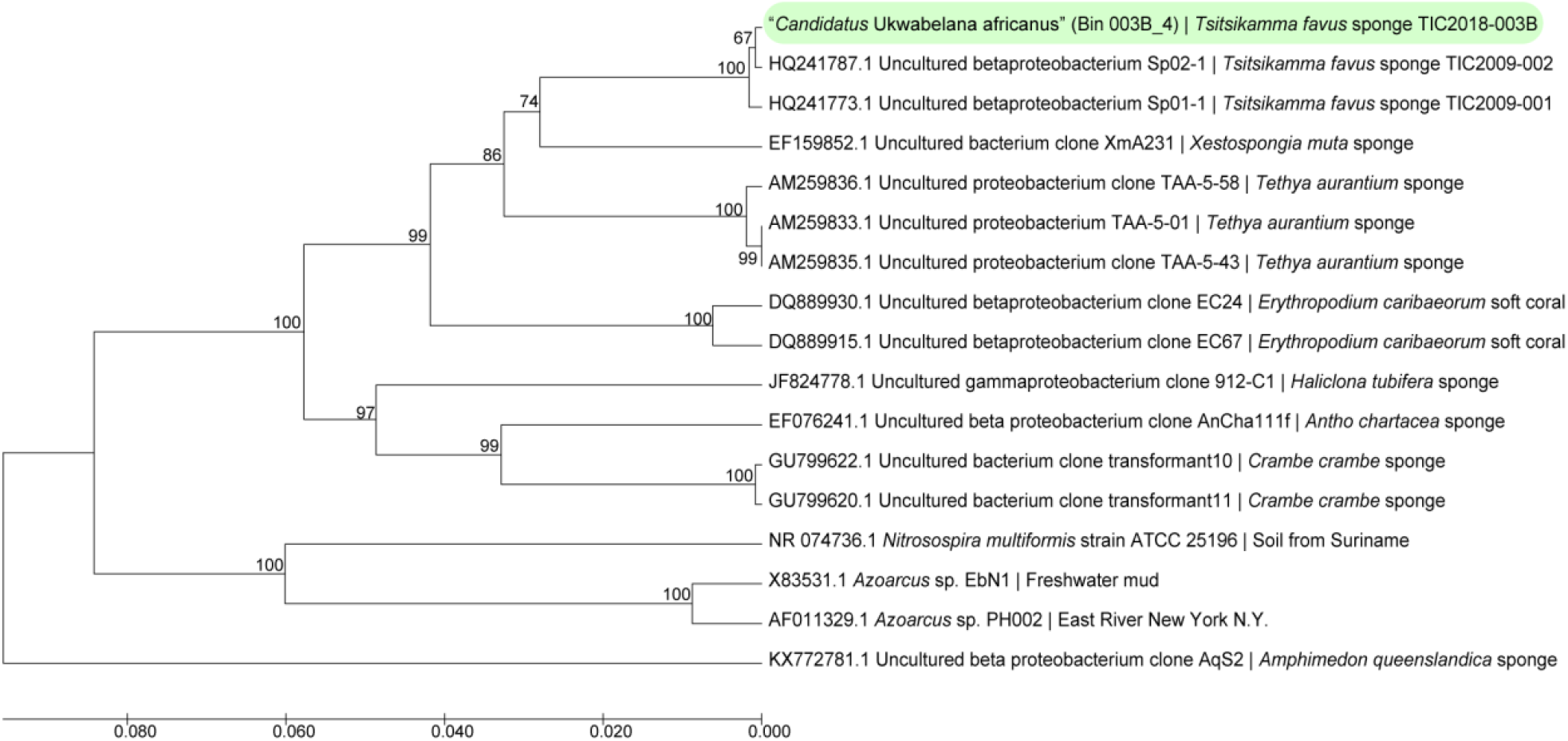
Phylogeny of *T. favus*-associated betaproteobacterium bin. The phylogenetic relationship between the putative betaproteobacteria Sp02-1 genome and closest relatives was based on 16S rRNA sequences and inferred using the UPGMA method. The percentage of replicate trees in which the associated taxa clustered together in the bootstrap test (10000 replicates) are shown next to the branches. The tree is drawn to scale, with branch lengths in the same units as those of the evolutionary distances used to infer the phylogenetic tree. The evolutionary distances were computed using the Maximum Composite Likelihood method and are in the units of the number of base substitutions per site. This analysis involved 17 16S rRNA gene sequences with a total of 1291 positions analyzed. All ambiguous positions were removed for each sequence pair (pairwise deletion option). Evolutionary analyses were conducted in MEGA X.

**Figure S3.**
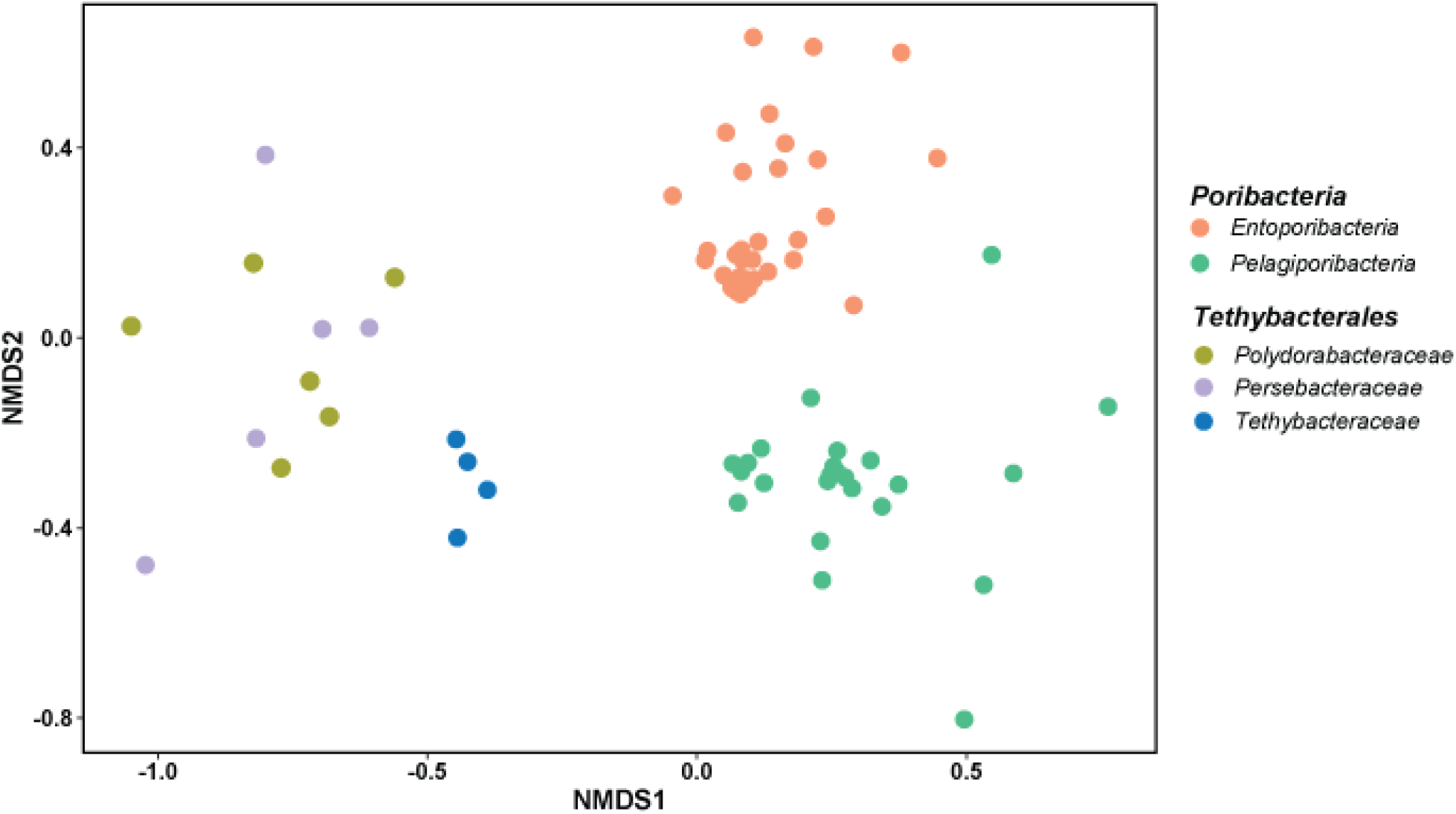
The potential functional ability of the three Tethybacterales families and the two lineages within the Poribacteria appear to be distinct. A Non-Metric Multi-Dimensional Scaling (NMDS) plot of the presence/absence metabolic counts from the Tethybacterales and Poribacteria calculated using Bray-Curtis distance.

